# Icariin induces MC3T3-E1 cell proliferation and differentiation via the BMP-2/Smads/Runx2 signal pathway

**DOI:** 10.1101/405787

**Authors:** Qing An, Bo Gou, Shaocheng Ma, En Lin Goh, GuoXiong Liu, Bikash Kumar Sah, Hong Cao

**Affiliations:** Department of Orthopedic Surgery, Renmin Hospital, Hubei University of Medicine, Shiyan 442000, Hubei, P.R. China; Mechanical Engineering Department, Faculty of Engineering, Imperial College London, London, SW7 2AZ, United Kingdom; Faculy of Medicine, Imperial College London, London, W6 8PR, United Kingdom

**Keywords:** Icariin, MC3T3-E1 cells, Osteoporosis, Signal pathway

## Abstract

Icariin, the main active ingredient of Epimedium, has played an important role in bone anabolism. However, the molecular mechanism for this effect was not convincingly reported yet. In this paper, the role of icariin on cell morphology, viability, cell cycling and the activity of alkaline phosphatase (ALP) were studied, and the molecular mechanism of icariin induced osteogenic effect was also investigated. Icariin with different concentrations (10, 20 and 40 ng/ml) was used to modify the pre-osteoblastic MC3T3-E1 cells for 48, 72 and 96 h, and the proliferation, morphology, and the cell cycle of the cells were evaluated by Cell Counting Kit-8 (CCK-8), microscopy and flow cytometry, respectively. Bone morphogenic protein-2 (BMP-2), bone morphogenic protein receptor-2 (BMPR-2), Smad4, Smadl/5/8 proteins expression levels were obtained by Western blotting and the expression levels of runt-related transcription factor 2 (Runx2) mRNA was examined by reverse transcription-polymerase chain reaction (RT-PCR). In this study, we found that icariin could promote the proliferation and differentiation of MC3T3-E1 cells in a dose - and time-dependent manner. Icariin could stimulate the expression of the BMP-2, BMPR-2, Smad4 and Smadl/5/8 proteins. Furthermore, icariin could upregulate the expression of Runx2 mRNA. These results showed that icariin played an important role in upregulating BMP-2 expression to activate the BMP-2/Smads/Runx2 signal pathway for increasing both the proliferation and differentiation of the MC3T3-E1 cells. However, the osteogenic effects of icariin can be suppressed by the BMP-2 antagonist (Noggin). In conclusion, we demonstrate that icariin is an osteoinductive factor that exerts its osteogenic effect by regulating the BMP-2/Smads/Runx2 signal pathway in MC3T3-E1 cells.

## Introduction

Aging-associated diseases have become a global challenge increasing the healthcare costs and lowering the quality of life. Osteoporosis, especially the postmenopausal osteoporosis (PMO) is a typical aging-associated disease and normally coexists with hypogonadism[1]. It becomes a significant issue with over 200 million people suffered worldwide[2]. Bisphosphonate is considered as a corner stone treatment for osteoporosis but recent research showed bisphosphonate may not significantly lower the fracture risk in a long treatment (> 5 years)[3, 4]. Alternatively, hormone replacement therapy (HRT) is an effective method to prevent postmenopausal osteoporosis[5]. However, the side effect of HRT is a major problem, such as the higher risk to cause breast cancer, heart disease, stroke and pulmonary embolism[6]. Therefore, it is urgent to find safe and effective drugs to mitigate and treat osteoporosis.

Icariin (C33H40O15; molecular weight, 676.67) is an important component in Herba epimedii, which has been used in Chinese herbal medicine to treat osteoporosis for centuries[7, 8]. Our previous study found that icariin was potentially a novel therapeutic ingredient for fracture recovery in postmenopausal osteoporosis[9]. In addition, numerous studies have shown that icariin has anti-osteoporosis effect by stimulating bone formation as well as suppressing bone resorption[10-12]. However, the molecular mechanism has not been fully elucidated. Cao et al.[13] and Yin et al.[14] suggested that icariin might exert its osteogenic effects through the production of BMP-2 in osteoblasts. Zhou et al.[15] reported that icariin played a critical role in differentiating MC3T3-E1 osteoblastic cells in vitro, possibly by promoting Smad1 and Smad5 production in osteoblasts. Furthermore, studies also demonstrated that icariin stimulated osteogenesis by promoting Runx2 expression in vitro[16, 17]. Taken together, we hypothesized that icariin could promote the expression of BMP-2 to activate the BMP-2/Smads/Runx2 signaling pathway to induce bone formation.

Therefore, we investigated the role of icariin on cell morphology, cell viability and alkaline phosphatase (ALP) viability. We also studied the effect of icariin on BMP-2, BMPR-2, Smad4 and Smadl/5/8 proteins and the Runx2 mRNA expression in MC3T3-E1 cells, which were involved in BMP-2/Smads/Runx2 signaling pathway.

## Materials and methods

### Reagents and Cell culture

Icariin was obtained from the National Institutes for Food and Drug Control (Beijing, China) with a purity of 99%. Icariin powder was dissolved in dimethylsulfoxide (DMSO; Sigma, St. Louis, MO, USA) and kept at −20 °C before use. The test solutions of icariin (0, 10, 20 and 40 ng/ml) were prepared from the stock solution by dilution with the culture medium. MC3T3-E1 cells were obtained from the Institute of Biochemistry and Cell Biology (Shanghai, China) and maintained as reported. Primary antibody BMP2 (ab14933), BMPR2 (ab96826), Smad4 (ab40759) were from Abcam (Cambridge, MA, USA) and primary antibody Smad1/5/8/9 (NB100-56443) were from Novus Biotechnology.

### Hematoxylin and eosin (H&E) staining

MC3T3-E1 cells were loaded onto 6-well plates (1×105 cells/ml) and cultured for 24 h with the standard conditions, then the culture medium was replaced with icariin solutions (0, 10, 20 and 40 ng/ml) and the cells were treated for 48, 72 and 96 h at each icariin concentration. For morphological observation, MC3T3-E1 cells were stained using hematoxylin and eosin (H&E) and observed using a microscope.

### Cell viability assay

The MC3T3-E1 cell viability was measured by Cell Counting Kit-8 kit (CCK-8) (Shanghai Yisheng Biotechnology Co. Ltd, Shanghai, China). MC3T3-E1 cells were loaded onto 96-well plates (1×105 cells/ml) then cultured and treated the same as in the H&E staining section. The viability of the MC3T3-E1 cells based on its absorbance, which was measured on microplate reader at 450 nm (BioTek Instruments, USA).

### Cell cycle assay

MC3T3-E1 cells were incubated with icariin solutions as in the H&E section for 72. The cells were digested by trypsin and fixed in 70% ethanol for 12 h at 4 °C. After the cells were incubated with RNase A (Invitrogen, Carlsbad, CA, USA) and propidium iodide (PI), the cell cycle assay was performed on flow cytometry.

### ALP activity assay

The ALP activity was measured using Alkaline Phosphatase Assay Kit (Yisheng Biotechnology Co. Ltd, Shanghai, China) on microplate reader at 48, 72 and 96 h of culture. Values of ALP activity were normalized based on the protein concentration, which was calculated from the bicinchoninic acid (BCA) protein assay.

### Western blot analysis

The MC3T3-E1 cells were cultured as in the H&E staining section. The cells were lysed with RIPA at 4 °C for 1 h and the then total protein was extracted as reported. Proteins were separated using 12% SDS-PAGE at a constant voltage (72 V), and electroblotted on polyvinylidene difluoride (PVDF) membranes at a constant-current (250 mA) for approximately 1 h. 5% non-fat milk/PBS solution were used to saturate the non-specific binding sites for 1 h. The membranes were incubated with BMP-2 (1:1,000), BMPR-2 (1:1,000), Smad4 (1:1,000) and Smad1/5/8 (1:200) primary antibodies at 4 °C overnight and then the blots were detected with secondary antibody at room temperature for 2 h with rocking. Finally, bands were quantified by enhanced chemiluminescence and normalized to β-actin concentration.

### RT-PCR

The cells were prepared and treated for 72 h as in the H&E staining section. The cells were lysed with TRIzol and the total RNA was harvested as reported. RevertAid(tm) First Strand cDNA Synthesis Kit (Fermentas, USA) was used to generate cDNA. The sequences of the PCR primers of Runx2 as follows: Forward sequence (5′ to 3′) AGC GGA CGA GGC AAC AGT TT and reverse sequence (5′ to 3′) CCT AAA TCA CTG AGG CGG TCA G. GAPDH forward sequence (5′ to 3′) CAA CTA CAT GGT TTA CAT GTT C and reverse sequence (5′ to 3′) GCC AGT GGA CTC CAC GAC. The PCR amplification reaction mixture contained 5 μl Light Cycler FastStart DNA Master SYBR Green I (Roche, Mannheim, Germany), 3 μl of cDNA template and 2 μl PCR primers. Amplification was performed as following: 94 °C for 5 min, 94 °C for 20 sec, 55 °C for 10 sec and 72 °C for 10 sec. The PCR products were probed by agarose gel electrophoresis and normalized based on the glyceraldehyde 3-phosphate dehydrogenase (GAPDH) concentration.

## Statistical analysis

Unpaired two-tailed Student’s test of the GraphPad Prism Version 6.00 analysis software (GraphPad Software, Inc., San Diego, CA) was used to analyze the experimental results. Results were presented as the mean ± standard deviation (SD). P < 0.05 was considered to indicate a statistically significant difference.

## Results

### Effect of icariin on cell morphology

Cell morphology of treated MC3T3-E1 was observed by H&E dying and microscopy. At 48 and 72 h, cells appeared polygonal and fusiformis and the nucleus was clearly visible in each group (Fig. 1-1, Fig. 1-2). At 96 h, cells were arranged closely. The cells appeared polygonal and scale-shaped and protruded more protrusions, which interconnected with each other (Fig. 1-3). No observable differences of MC3T3-E1 morphology for each group were found. The results indicate that the icariin has no detectable effect on the morphology of MC3T3-E1 cells.

**Figure 1-1.**
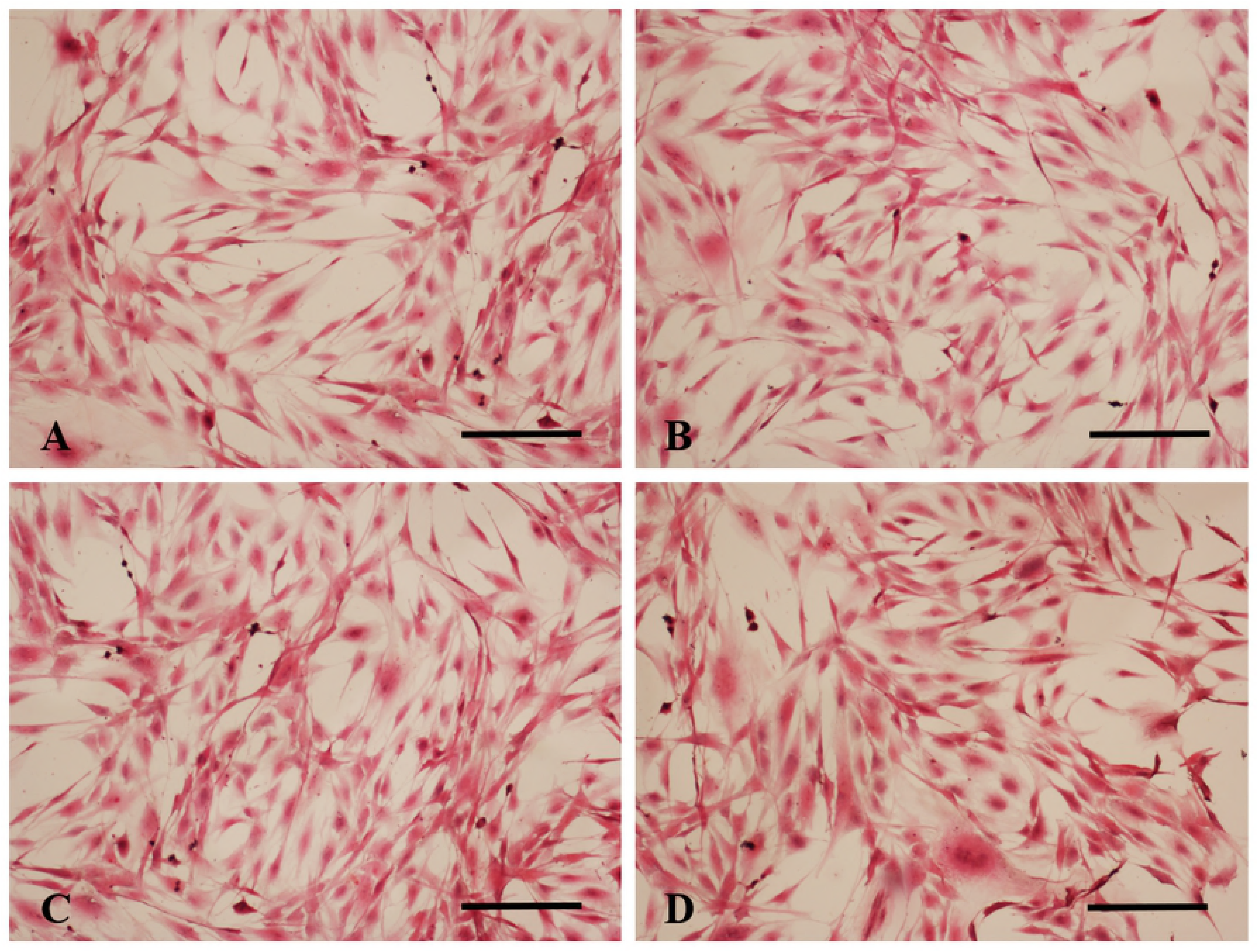
Morphology of MC3T3-E1 cells treated with (A) 0, (B) 10, (C) 20 and (D) 40 ng/ml icariin for 48 h. (Hematoxylin and Eosin staining. Bars indicate 100 μm).

**Figure 1-2.**
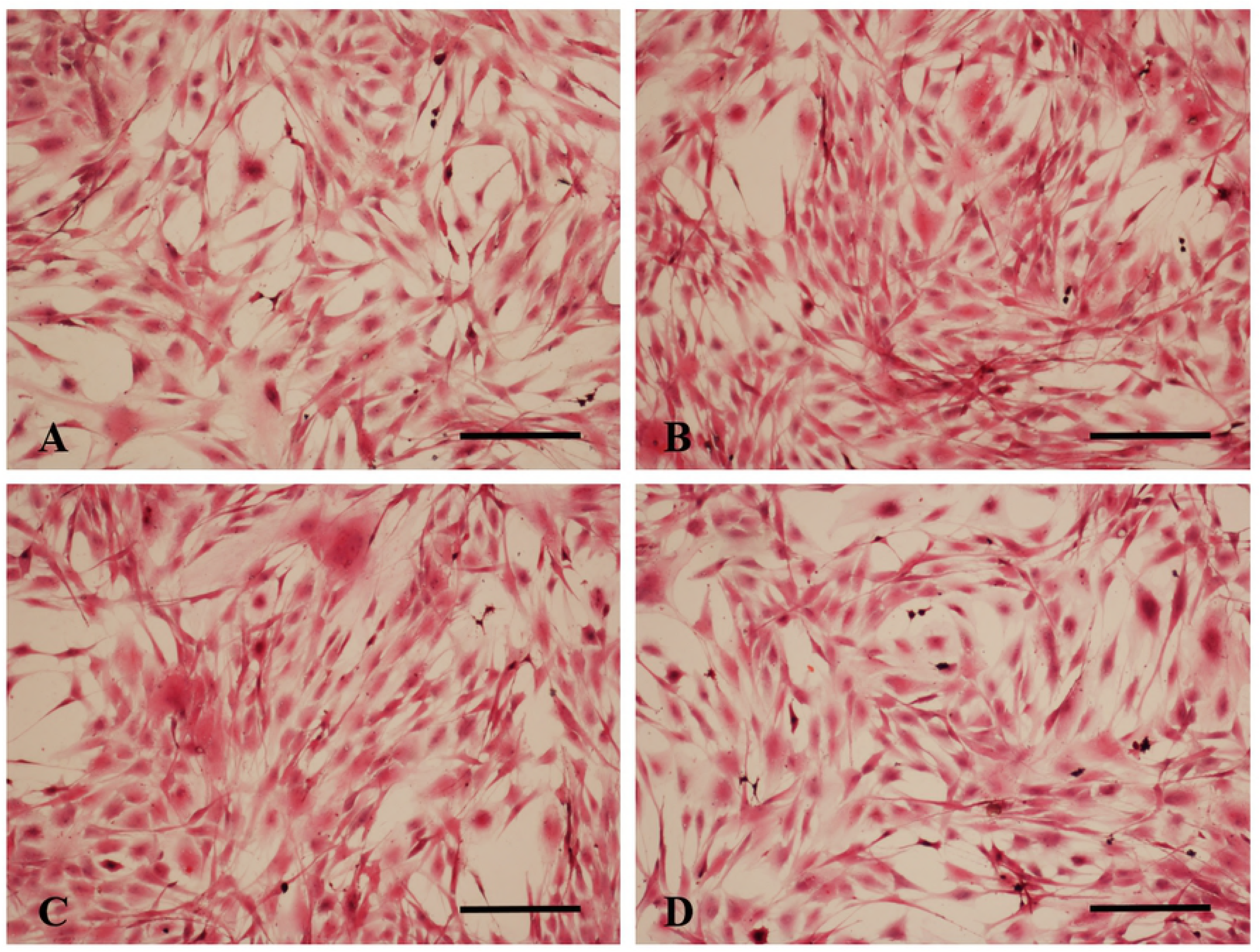
Morphology of MC3T3-E1 cells treated with (A) 0, (B) 10, (C) 20 and (D) 40 ng/ml icariin for 72 h. (Hematoxylin and Eosin staining. Bars indicate 100 μm).

**Figure 1-3.**
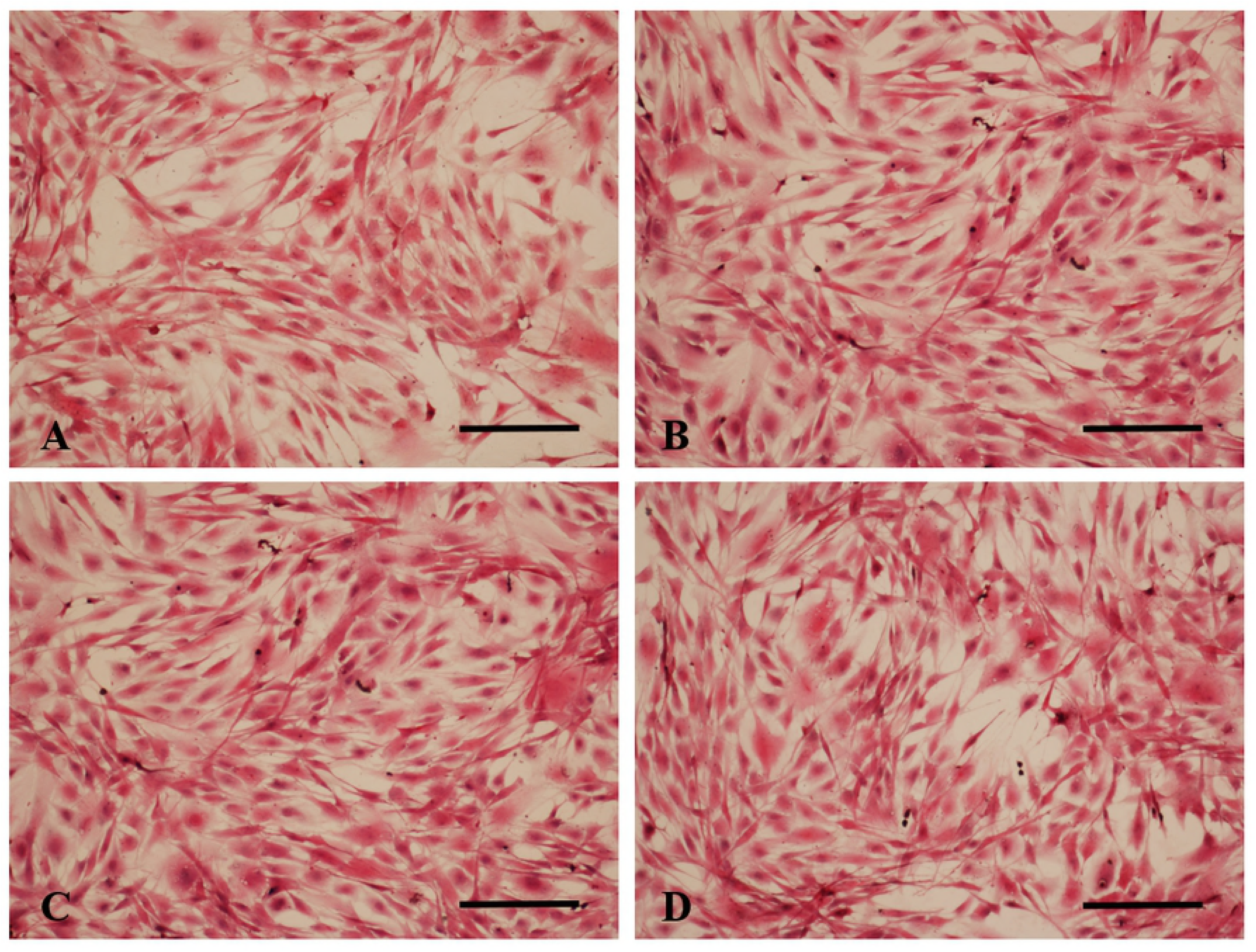
Morphology of MC3T3-E1 cells treated with (A) 0, (B) 10, (C) 20 and (D) 40 ng/ml icariin for 96 h. (Hematoxylin and Eosin staining. Bars indicate 100 μm).

### Effect of icariin on cell viability

The cell viability treated with 10 and 20 ng/ml of icariin was significantly increased compared to the control without icariin (P < 0.01) (Fig. 2A). The maximal cell viability was the cell treated with the 20 ng/ml of Icariin. Cell viability with 20 ng/ml of icariin at 48, 72 and 96 h was significantly higher than that at 24 h and peaked on 96 h. The results demonstrate that icariin could increase MC3T3-E1 cells proliferation according to the dose and time.

**Figure 2.**
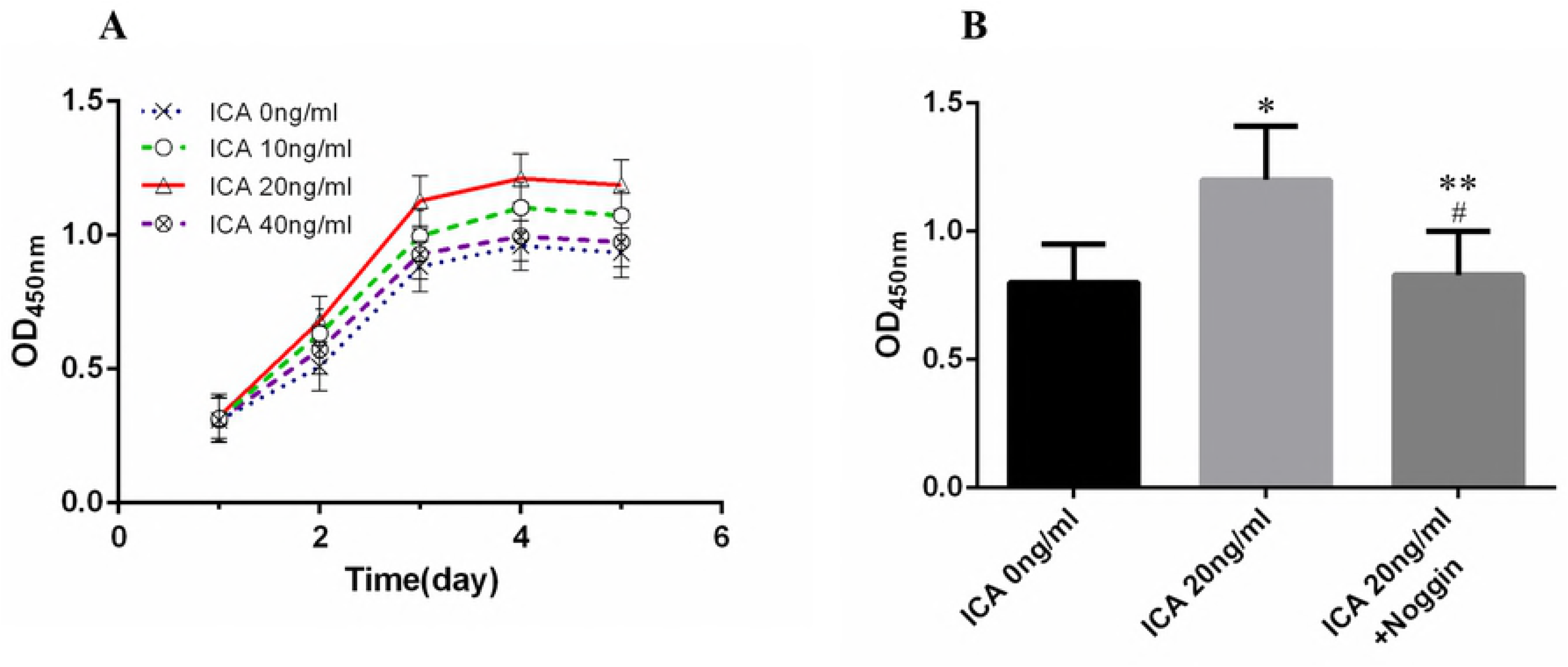
Effect of Icariin on cell viability in MC3T3-E1 cells. (A) The proliferation of MC3T3-E1 cells was detected by CCK-8 assay after treatment with different concentrations of icariin (0, 10, 20 and 40 ng/ml) for 48, 72 and 96 h. (B) MC3T3-E1 cells were cultured with 0 and 20 ng/ml icariin in the presence or absence of Noggin (100 ng/ml) for 72 h and then detected by CCK-8 assay. Each value is the mean ± SD, \**P* < 0.01; #*P* > 0.05, versus the control group; \*\**P* < 0.01, (Icariin 20 ng/ml) group versus (Icariin 20 ng/ml + Noggin) group.

### Effect of icariin on cell cycle division

When MC3T3-E1 cells were cultured with icariin for 72 h, the group with 10 and 20 ng/ml of icariin showed increased ratios of S-phase cells and decreased ratios of G0/G1-phase cells compared to control group (0 ng/ml) (Fig. 3-1). These data demonstrated that icariin could increase MC3T3-E1 cell proliferation by stimulatingthe cell division.

**Figure 3-1.**
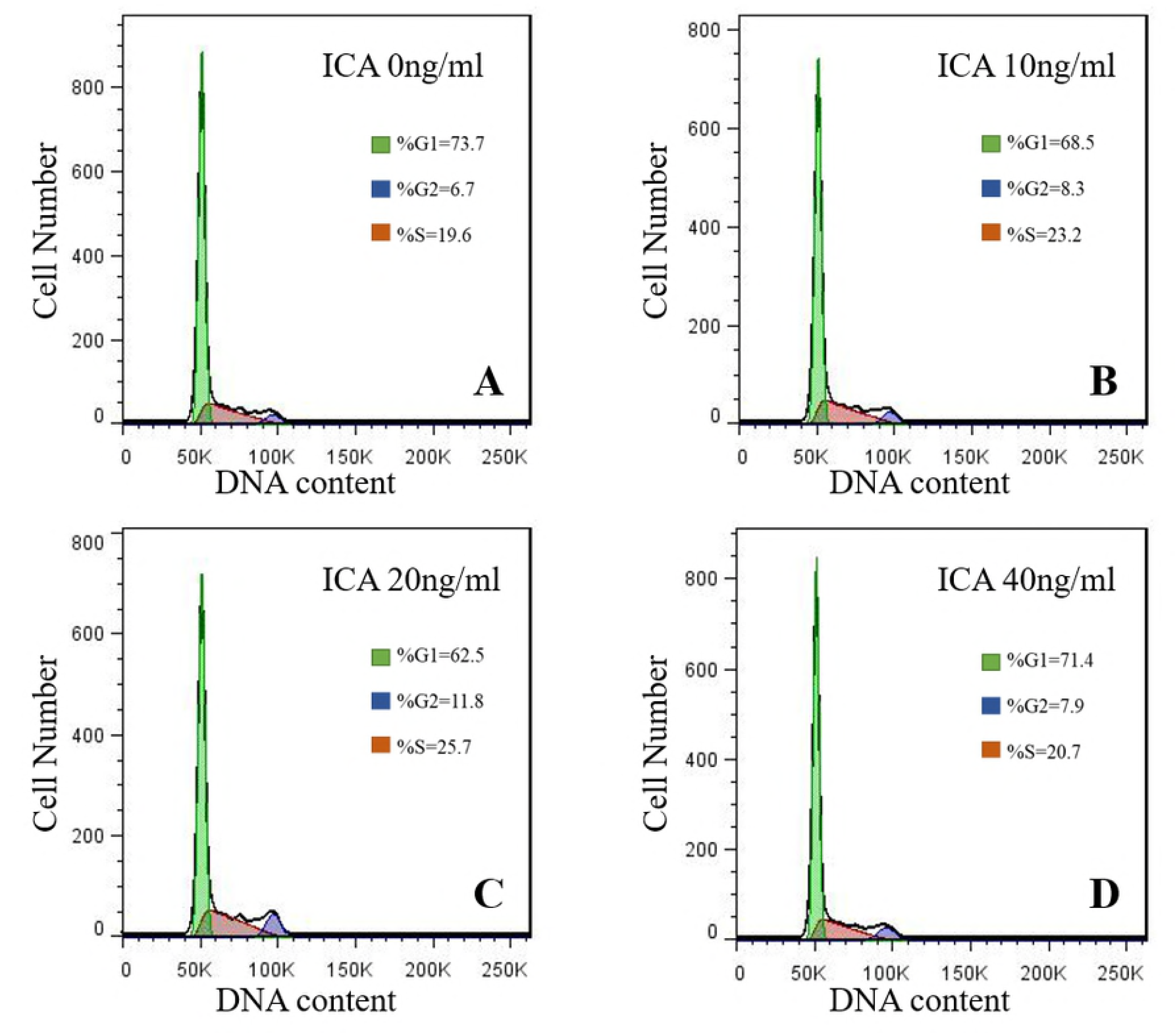
Effect of icariin on MC3T3-E1 cell cycle was detected by flow cytometry analysis. MC3T3-E1 cells treated with (A) 0, (B) 10, (C) 20 and (D) 40 ng/ml concentrations of icariin for 72 h.

**Figure S3-2.**
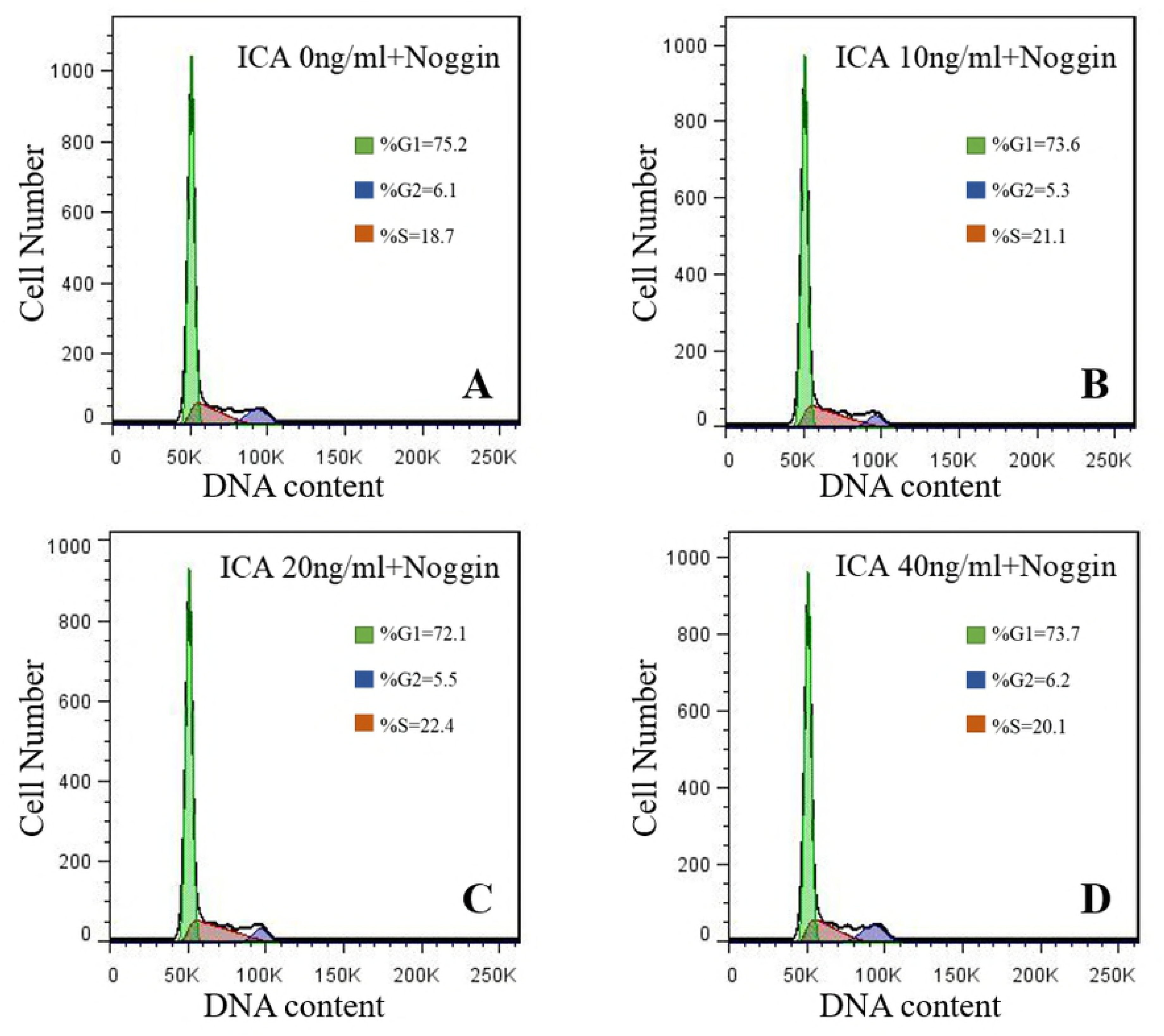
Effect of Noggin on cell cycle induced by icariin in MC3T3-E1 cells. The MC3T3-E1 cells were co-cultured with different concentrations of icariin (0, 10, 20 and 40 ng/ml) and Noggin (100 ng/ml) and were examined by flow cytometry analysis at 72 h.

### Effect of icariin on ALP activity

The ALP activity among different groups (0, 10, 20 and 40 ng/ml) showed no obvious differences at 24 h. Whereas at 48, 72 and 96 h, the ALP activity increased significantly in 10 and 20 ng/ml icariin groups compared to the control (P < 0.01) (Fig. 4A). ALP activity peaked on 96 h in 20 ng/ml icariin group and declined thereafter. As ALP is a maker of early osteoblast differentiation, these results demonstrate that icariin could promote MC3T3-E1 cells differentiation based on the dose and incubation time.

**Figure 4.**
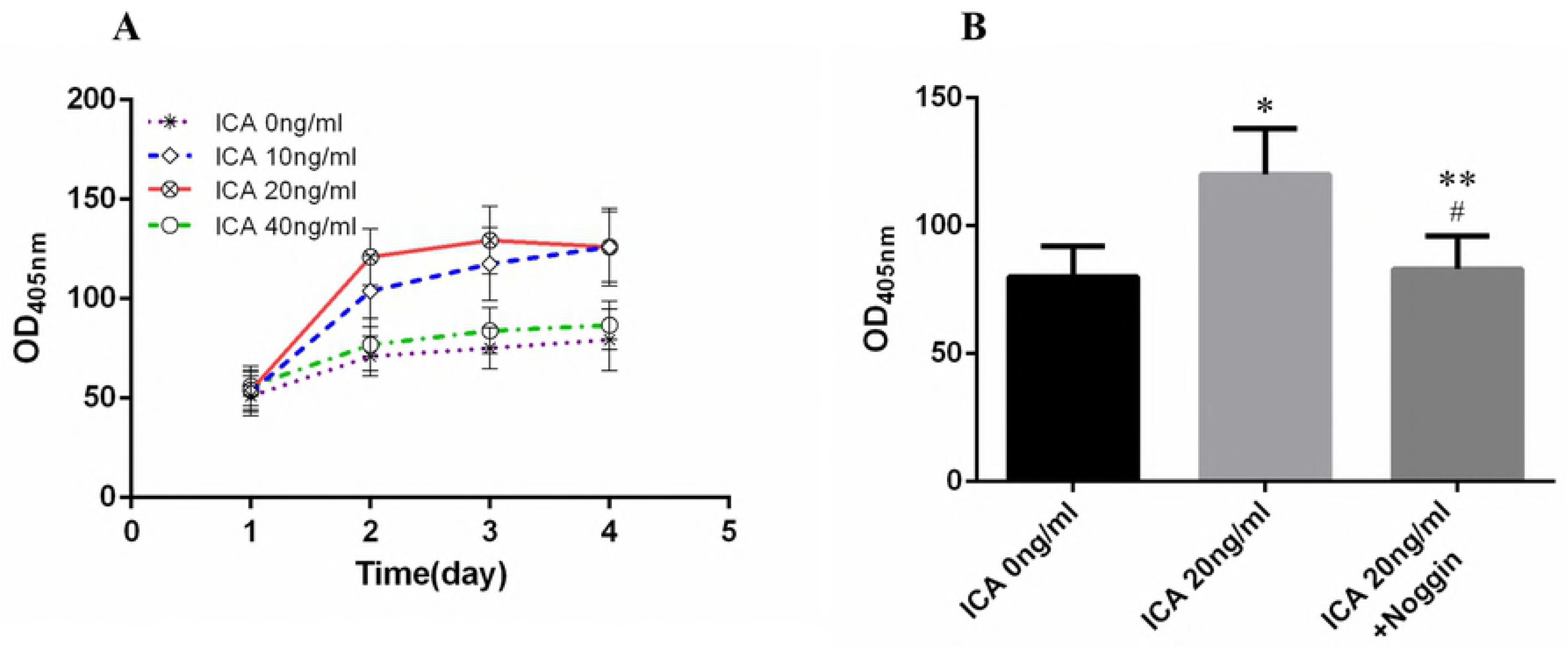
Effect of icariin on alkaline phosphatase (ALP) activities were measured at 405 nm on microplate reader. (A) The MC3T3-E1 cells were treated with different concentrations of icariin (0, 10, 20 and 40 ng/ml) for 48, 72 and 96 h. (B) MC3T3-E1 cells were cultured for 72 h in 0 and 20 ng/ml icariin groups in the presence or absence of Noggin (100 ng/mL). Each value is the mean ± SD, \**P* < 0.01; #*P* > 0.05, versus the control group; \*\**P* < 0.01, (Icariin 20 ng/ml) group versus (Icariin 20 ng/ml + Noggin) group.

### Effect of icariin on BMP-2, BMP-2R, Smad1/5/8 and Smad4 proteins expression levels

The effect of icariin concentration on BMP-2, BMP-2R, Smad1/5/8 and Smad4 protein expression levels was examined by Western blotting on 48, 72 and 96 h (Fig. 5-1A). 10 and 20 ng/ml icariin significantly increased BMP-2, Smad4, Smad1/5/8, and BMP-2R expression in the MC3T3-E1 cells compared to the control (0 ng/ml) (P < 0.01) (Figs. 5-1B, 5-1C, 5-1D, and 5-1E respectively). These results showed that icariin could promote BMP-2, BMP-2R, Smad1/5/8 and Smad4 protein expression, suggesting that icariin might promote MC3T3-E1 cells proliferation and differentiation via its actions on BMP-2/Smads signal pathway.

**Figure 5-1.**
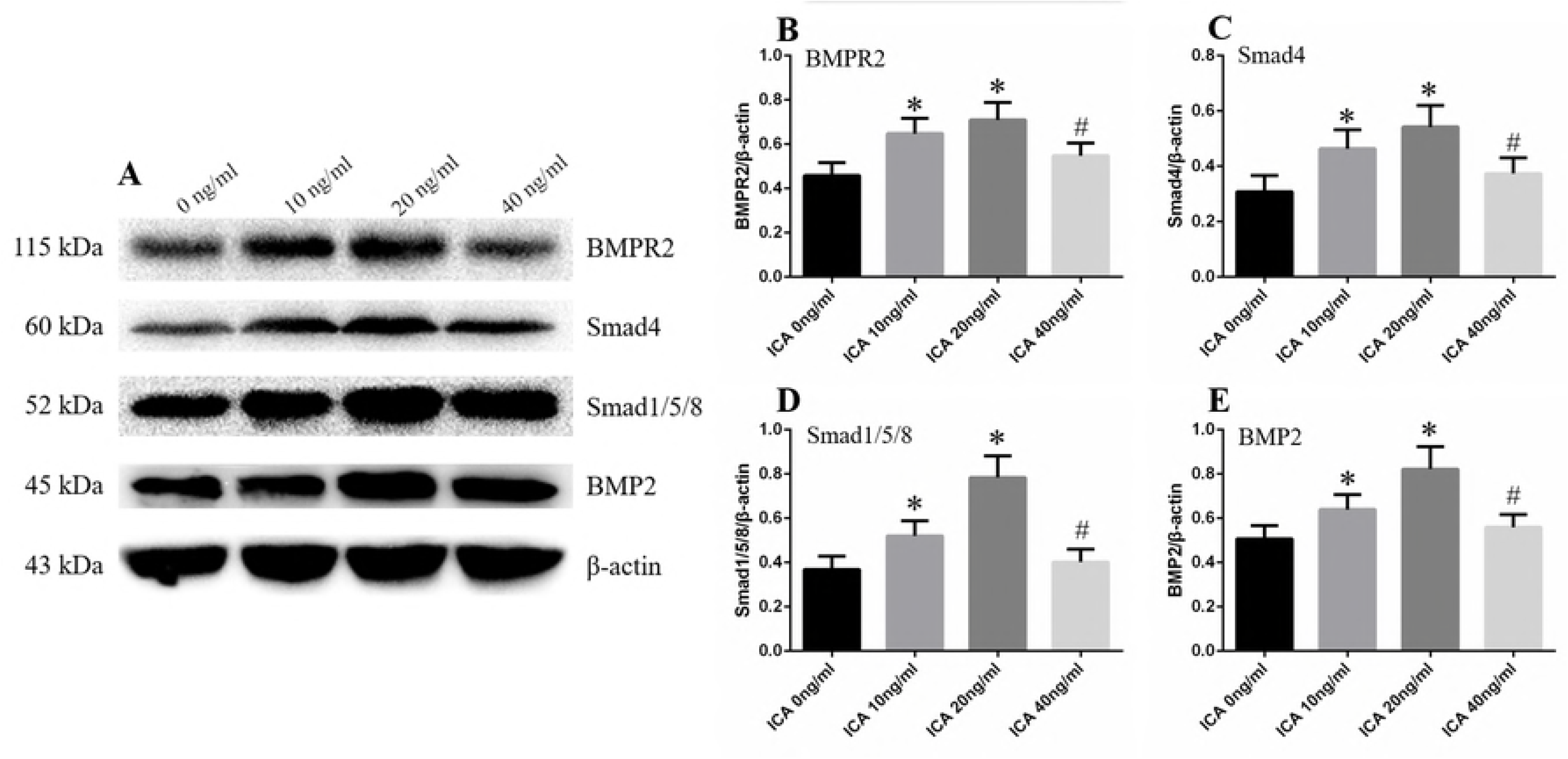
Effect of icariin on BMP-2, BMP-2R, Smad1/5/8 and Smad4 proteins expression in MC3T3-E1 cells. The MC3T3-E1 cells were treated with different concentrations of icariin (0, 10, 20 and 40 ng/ml) for 72 h. All the protein was isolated and (A) western blot analysis was performed to measure the protein expression levels of (B) BMPR2, (C) Smad4, (D) Smad1/5/8 and (E) BMP2, which were normalized on the basis of β-actin concentration. Each value is the mean ± SD, \**P* < 0.01; #*P* > 0.05, versus the control group (Icariin 0 ng/ml).

**Figure 5-2.**
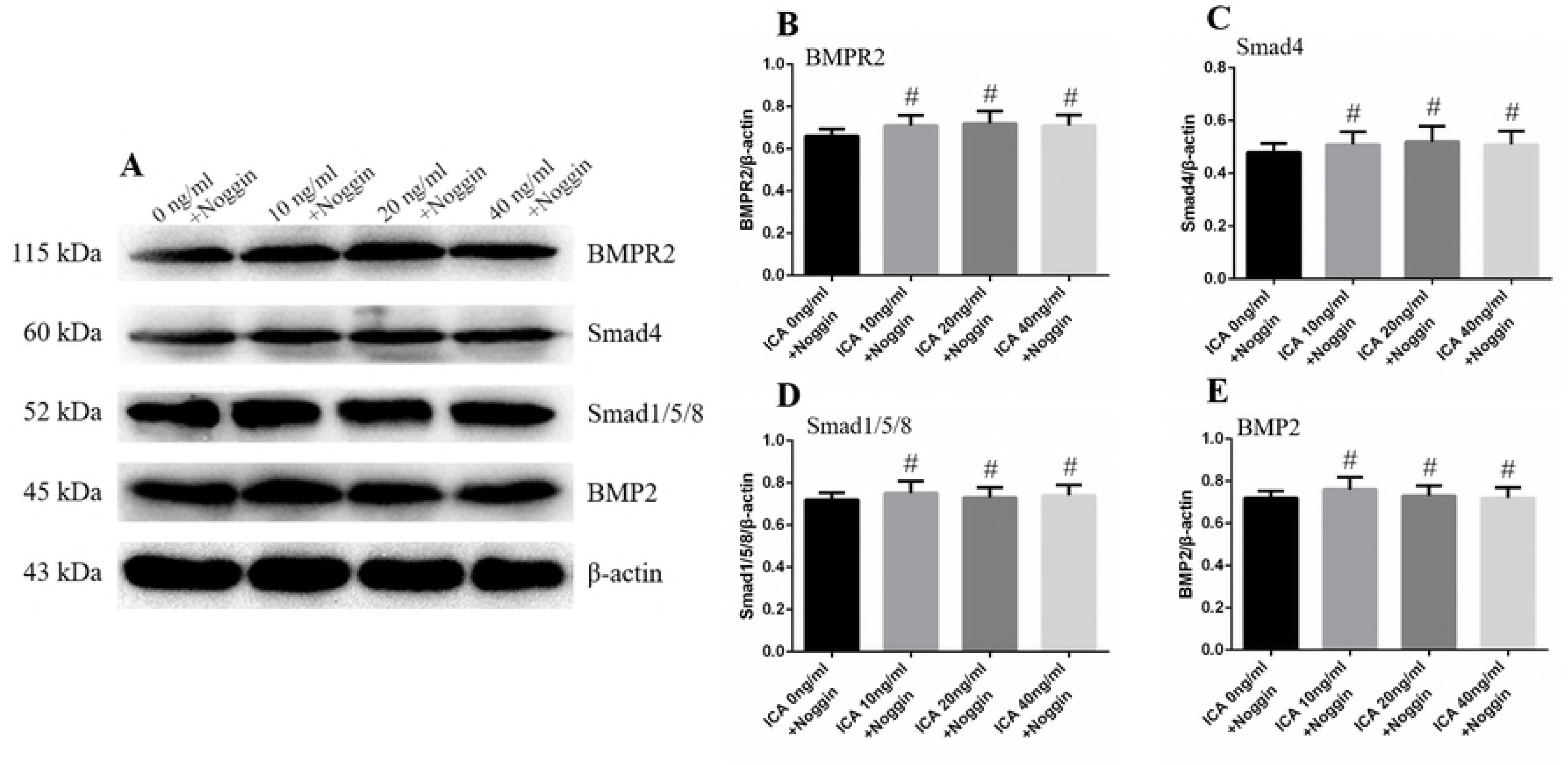
Effect of Noggin on BMP-2, BMP-2R, Smad1/5/8 and Smad4 proteins expression induced by icariin in MC3T3-E1 cells. The MC3T3-E1 cells were co-cultured with different concentrations of icariin (0, 10, 20 and 40 ng/ml) and Noggin (100 ng/mL) for 72 h. All the protein was isolated and (A) western blot analysis was performed to measure the protein expression levels of (B) BMPR2, (C) Smad4, (D) Smad1/5/8 and (E) BMP2, which were normalized on the basis of β-actin concentration. Each value is the mean ± SD, #*P* > 0.05, versus the control group (Icariin 0 ng/ml).

### Effect of icariin on Runx2 mRNA expression

When MC3T3-E1 cells were incubated with icariin for 72 h, RT-PCR showed that Runx2 genes were highly up-regulated in the 10 and 20 ng/ml group. The Runx2 mRNA expression level in the 40 ng/ml icariin group was slightly increased compared to the control (0 ng/ml), while lower than that of the 10 and 20 ng/ml groups (P < 0.05) (Fig. 6-1). This result indicated that certain concentrations of icariin could significantly increase the Runx2 mRNA expression in MC3T3-E1 cells.

**Figure 6-1.**
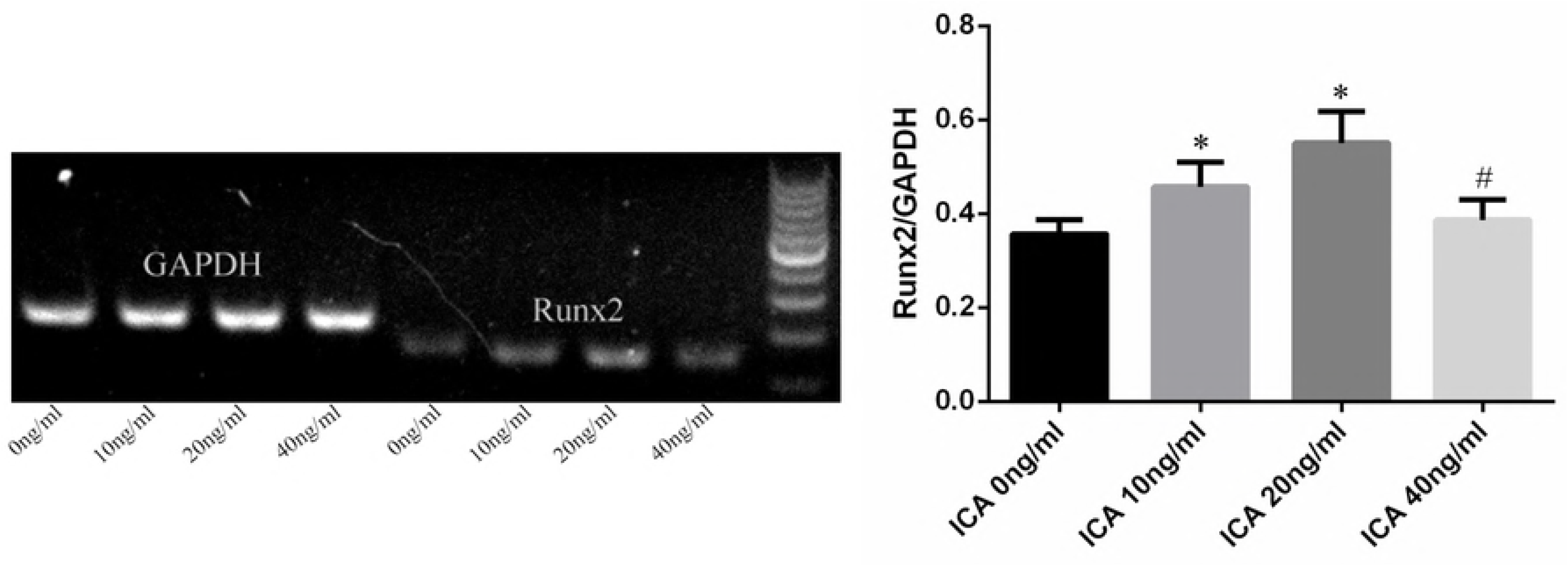
Effect of icariin on Runx2 mRNA expression in MC3T3-E1 cells. The MC3T3-E1 cells were treated with different concentrations of icariin (0, 10, 20 and 40 ng/ml) for 72 h. All the RNA was isolated and real-time PCR was performed to measure the mRNA expression levels of Runx2, which were normalized on the basis of GAPDH concentration. Each value is the mean ± SD, \**P* < 0.01; #*P* > 0.05, versus the control group (Icariin 0 ng/ml).

**Figure 6-2.**
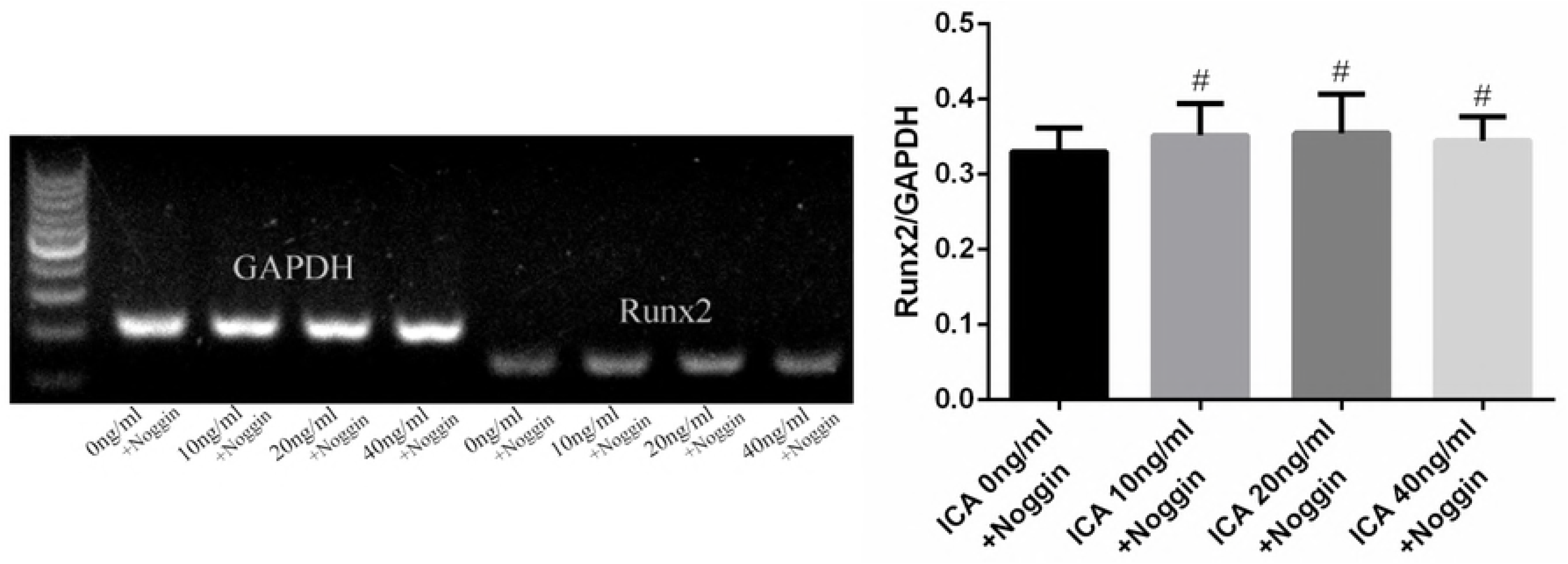
Effect of Noggin on the osteogenic effect of icariin in MC3T3-E1 cells. The MC3T3-E1 cells were co-cultured with different concentrations of icariin (0, 10, 20 and 40 ng/ml) and Noggin (100 ng/mL) for 72 h. All the RNA was isolated and real-time PCR was performed to measure the mRNA expression levels of Runx2, which were normalized on the basis of GAPDH concentration. Each value is the mean ± SD, \**P* < 0.01; #*P* > 0.05, versus the control group (Icariin 0 ng/ml).

### Noggin could block the osteoblast proliferation and differentiation induced by icariin

To further explore the role of BMP-2 in proliferation and differentiation induced by icariin, MC3T3-E1 cells were co-cultured with icariin (0, 10, 20 and 40 ng/ml) and BMP-2 antagonist (Noggin; 100 ng/ml) (Sigma Chemical, USA). Noggin significantly inhibited the icariin induced cell proliferation (Fig. 2B), cell cycle division (Fig. 3-2) and ALP activity (Fig. 4B), and suppressed protein expression levels of the BMP-2 (Fig. 5-2E), BMP-2R (Fig. 5-2B), Smad1/5/8 (Fig. 5-2D) and Smad4 (Fig. 5-2C) proteins and the gene expression levels of Runx2 induced by icariin (Fig. 6-2). These data demonstrate that icariin could induce MC3T3-E1 cells proliferation and differentiation via the BMP-2/Smads/Runx2 signal pathway.

## Discussion

Icariin is a major flavonoid in Herba Epimedii, which has an osteogenic effect[18-20]. Studies have reported that icariin could increase the proliferation and differentiation of pre-osteoblasts, osteoblasts and rat bone marrow stromal cells[21, 22]. Song et al.[23] reported that icariin could increase MC3T3-E1 cell proliferation and reduce cell apoptosis. Both of the cell proliferation and apoptosis are associated with increased mRNA levels of positive regulators of cell cycle gene Cyclin E and proliferating cell nuclear antigen (PCNA). In the same report, Song et al.[23] also showed that icariin could decrease mRNA level of negative regulator gene, Cyclin-dependent kinase 4 inhibitor B (Cdkn2B), and reduce caspase-3 activity. Our results also showed that icariin could increase the MC3T3-E1 cells proliferation by stimulating cell cycle division.

ALP, as a marker of early osteoblast differentiation, began to appear in the extracellular matrix synthesis phase and reached its peak in the mineralized nodule formation phase[24]. In this study, MC3T3-E1 cells cultured in the 10 and 20 ng/ml of icariin exhibited the significantly increase in ALP activity (P < 0.01). Our results indicate that icariin has osteogenesis function by stimulating osteoblasts differentiation in vitro.

Bone Morphogenetic Proteins (BMPs) are in the transforming growth factor-β (TGF-β) superfamily, which is important in the regulation of bone induction, maintenance and repair[25]. BMP-2 has been considered to be the most active osteogenesis factor in TGF-β superfamily. Studies showed that BMP-2 could strongly promote the proliferation and differentiation of pre-osteoblasts, osteoblasts and bone marrow stromal cells[26, 27]. BMP-2 binds to the transmembrane receptor (BMPR) to activate the heterotetrameric serine/threonine kinase. The activated receptor then phosphorylates the regulatory Smadl/5/8, which interacts with the common mediator Smad (Smad4) and forms a heteromeric complex[27]. The complex then relocates into the nucleus, thereby regulating the transcription of the target gene Runx2, which is a multifunctional transcription factor regulating the development of bone by promoting the differentiation of chondrocytes and osteoblasts[28]. This study suggested that icariin not only increased the expression levels of BMP-2 the same way as our previous study, but also promoted the BMPR-2, Smad4, Smadl/5/8 proteins which up-regulated the gene expression levels of Runx2 based on the dose. These results indicate that icariin might up-regulate the expression levels of BMP-2 to activate the BMP-2/Smads/Runx2 signal pathway, which regulates the MC3T3-E1 cell proliferation and differentiation. Continued studies are necessary to confirm this.

The changes of indexes mentioned above were observed when Noggin, a BMP antagonist[29], was added to MC3T3-E1 cells. It showed that Noggin blocked MC3T3-E1 cells proliferation and ALP activity induced by icariin. In addition, Noggin inhibited the expression levels of BMP-2, BMPR-2, Smad4, Smadl/5/8 proteins and Runx2 mRNA, which were involved in the BMP-2/Smads/Runx2 signal pathway induced by icariin. These results indicate that Noggin is capable of blocking the production of BMP-2 to inhibit the BMP-2/Smads/Runx2 signal pathway, which involves in the role of osteogenic effects induced by icariin. Therefore, it can be demonstrated that icariin is an anabolic agent that takes its osteogenic effect through the regulation of the BMP-2/Smads/Runx2 signal pathway in MC3T3-E1 cells. This study supports that icariin is a promising agent for the treatment and mitigation of osteoporosis.

Acknowledgments
This study was supported by the National Natural Science Foundation of China (No. 81602867), Hubei Provincial Department of Education (No. WJ2015Q042), and Hubei Provincial science and Technology Department funded projects (No. 2018CFB524).

## References

1. Hagino H, Soen S, Sugimoto T, Endo N, Okazaki R, Tanaka K, Nakamura T (2018) Changes in quality of life in patients with postmenopausal osteoporosis receiving weekly bisphosphonate treatment: a 2-year multicenter study in Japan. J BONE MINER METAB. https://doi.org/10.1007/s00774-018-0914-3

2. Johnell O, Kanis JA (2006) An estimate of the worldwide prevalence and disability associated with osteoporotic fractures. Osteoporos Int 17:1726–1733. https://doi.org/10.1007/s00198-006-0172-4

3. Ma S, Goh EL, Jin A, Bhattacharya R, Boughton OR, Patel B, Karunaratne A, Vo NT, Atwood R, Cobb JP et al. (2017) Long-term effects of bisphosphonate therapy: perforations, microcracks and mechanical properties. Sci Rep 7:43399. https://doi.org/10.1038/srep43399

4. Vasikaran SD (2009) Association of low-energy femoral fractures with prolonged bisphosphonate use: a case--control study. Osteoporos Int 20:1457–1458. https://doi.org/10.1007/s00198-009-0955-5

5. Grossman DC, Curry SJ, Owens DK, Barry MJ, Davidson KW, Doubeni CA, Epling JJ, Kemper AR, Krist AH, Kurth AE et al. (2017) Hormone Therapy for the Primary Prevention of Chronic Conditions in Postmenopausal Women: US Preventive Services Task Force Recommendation Statement. JAMA 318:2224– 2233. https://doi.org/10.1001/jama.2017.18261

6. Azam S, Lange T, Huynh S, Aro AR, von Euler-Chelpin M, Vejborg I, Tjonneland A, Lynge E, Andersen ZJ (2018) Hormone replacement therapy, mammographic density, and breast cancer risk: a cohort study. Cancer Causes Control 29:495–505. https://doi.org/10.1007/s10552-018-1033-0

7. Li C, Li Q, Mei Q, Lu T (2015) Pharmacological effects and pharmacokinetic properties of icariin, the major bioactive component in Herba Epimedii. LIFE SCI 126:57–68. https://doi.org/10.1016/j.lfs.2015.01.006

8. Kapoor S (2013) Icariin and its emerging role in the treatment of osteoporosis. Chin Med J (Engl) 126:400.

9. Cao H, Zhang Y, Qian W, Guo XP, Sun C, Zhang L, Cheng XH (2017) Effect of icariin on fracture healing in an ovariectomized rat model of osteoporosis. Exp Ther Med 13:2399–2404. https://doi.org/10.3892/etm.2017.4233

10. Luo Z, Liu M, Sun L, Rui F (2015) Icariin recovers the osteogenic differentiation and bone formation of bone marrow stromal cells from a rat model of estrogen deficiency-induced osteoporosis. Mol Med Rep 12:382–388. https://doi.org/10.3892/mmr.2015.3369

11. Zhang S, Feng P, Mo G, Li D, Li Y, Mo L, Yang Z, Liang (2017) Icariin influences adipogenic differentiation of stem cells affected by osteoblast-osteoclast co-culture and clinical research adipogenic. Biomed Pharmacother 88:436–442. https://doi.org/10.1016/j.biopha.2017.01.050

12. Zhang D, Zhang J, Fong C, Yao X, Yang M (2012) Herba epimedii flavonoids suppress osteoclastic differentiation and bone resorption by inducing G2/M arrest and apoptosis. Biochimie 94:2514–2522. https://doi.org/10.1016/j.biochi.2012.06.033

13. Cao H, Ke Y, Zhang Y, Zhang CJ, Qian W, Zhang GL (2012) Icariin stimulates MC3T3-E1 cell proliferation and differentiation through up-regulation of bone morphogenetic protein-2. Int J Mol Med 29:435–439. https://doi.org/10.3892/ijmm.2011.845

14. Yin XX, Chen ZQ, Liu ZJ, Ma QJ, Dang GT (2007) Icariine stimulates proliferation and differentiation of human osteoblasts by increasing production of bone morphogenetic protein 2. Chin Med J (Engl) 120:204–210.

15. Zhou H, Wang S, Xue Y, Shi N (2014) Regulation of the levels of Smad1 and Smad5 in MC3T3-E1 cells by Icariine in vitro. MOL MED REP 9:590–594. https://doi.org/10.3892/mmr.2013.1837

16. Liang W, Lin M, Li X, Li C, Gao B, Gan H, Yang Z, Lin X, Liao L, Yang M (2012) Icariin promotes bone formation via the BMP-2/Smad4 signal transduction pathway in the hFOB 1.19 human osteoblastic cell line. Int J Mol Med 30:889– 895. https://doi.org/10.3892/ijmm.2012.1079

17. Hsieh TP, Sheu SY, Sun JS, Chen MH, Liu MH (2010) Icariin isolated from Epimedium pubescens regulates osteoblasts anabolism through BMP-2, SMAD4, and Cbfa1 expression. Phytomedicine 17:414–423. https://doi.org/10.1016/j.phymed.2009.08.007

18. Tang D, Ju C, Liu Y, Xu F, Wang Z, Wang D (2018) Therapeutic effect of icariin combined with stem cells on postmenopausal osteoporosis in rats. J BONE MINER METAB 36:180–188. https://doi.org/10.1007/s00774-017-0831-x

19. Xu JH, Yao M, Ye J, Wang GD, Wang J, Cui XJ, Mo W (2016) Bone mass improved effect of icariin for postmenopausal osteoporosis in ovariectomy-induced rats: a meta-analysis and systematic review. Menopause 23:1152–1157. https://doi.org/10.1097/GME.0000000000000673

20. Fan JJ, Cao LG, Wu T, Wang DX, Jin D, Jiang S, Zhang ZY, Bi L, Pei GX (2011) The dose-effect of icariin on the proliferation and osteogenic differentiation of human bone mesenchymal stem cells. Molecules 16:10123–10133. https://doi.org/10.3390/molecules161210123

21. Chen KM, Ge BF, Ma HP, Liu XY, Bai MH, Wang Y (2005) Icariin, a flavonoid from the herb Epimedium enhances the osteogenic differentiation of rat primary bone marrow stromal cells. Pharmazie 60:939–942.

22. Ma XN, Zhou J, Ge BF, Zhen P, Ma HP, Shi WG, Cheng K, Xian CJ, Chen KM (2013) Icariin induces osteoblast differentiation and mineralization without dexamethasone in vitro. Planta Med 79:1501–1508. https://doi.org/10.1055/s- 0033-1350802

23. Song L, Zhao J, Zhang X, Li H, Zhou Y (2013) Icariin induces osteoblast proliferation, differentiation and mineralization through estrogen receptor-mediated ERK and JNK signal activation. Eur J Pharmacol 714:15–22. https://doi.org/10.1016/j.ejphar.2013.05.039

24. Witkowska-Sedek E, Stelmaszczyk-Emmel A, Majcher A, Demkow U, Pyrzak B (2018) The relationship between alkaline phosphatase and bone alkaline phosphatase activity and the growth hormone/insulin-like growth factor-1 axis and vitamin D status in children with growth hormone deficiency. Acta Biochim Pol. https://doi.org/10.18388/abp.2017_2541

25. Li Y, Hu W, Han G, Lu W, Jia D, Hu M, Wang D (2018) Involvement of bone morphogenetic protein-related pathways in the effect of aucubin on the promotion of osteoblast differentiation in MG63cells. Chem Biol Interact 283:51–58. https://doi.org/10.1016/j.cbi.2018.02.005

26. Egashira K, Sumita Y, Zhong W, I T, Ohba S, Nagai K, Asahina I (2018) Bone marrow concentrate promotes bone regeneration with a suboptimal-dose of rhBMP-2. PLoS ONE 13:e191099. https://doi.org/10.1371/journal.pone.0191099

27. Antebi YE, Linton JM, Klumpe H, Bintu B, Gong M, Su C, McCardell R, Elowitz MB (2017) Combinatorial Signal Perception in the BMP Pathway. CELL 170:1184–1196. https://doi.org/10.1016/j.cell.2017.08.015

28. Perez-Campo FM, Santurtun A, Garcia-Ibarbia C, Pascual MA, Valero C, Garces C, Sanudo C, Zarrabeitia MT, Riancho JA (2016) Osterix and RUNX2 are Transcriptional Regulators of Sclerostin in Human Bone. Calcif Tissue Int 99:302– 309. https://doi.org/10.1007/s00223-016-0144-4

29. Khattab HM, Kubota S, Takigawa M, Kuboki T, Sebald W (2018) The BMP-2 mutant L51P: a BMP receptor IA binding-deficient inhibitor of noggin. J Bone Miner Metab. https://doi.org/10.1007/s00774-018-0925-0

